# How a speaker herds the audience: Multi-brain neural convergence over time during naturalistic storytelling

**DOI:** 10.1101/2023.10.10.561803

**Authors:** Claire H. C. Chang, Samuel A. Nastase, Uri Hasson

## Abstract

Storytelling—an ancient way for humans to share individual experiences with others—has been found to induce neural synchronization among listeners. In our exploration of the dynamic fluctuations in listener-listener (LL) coupling throughout stories, we uncover a significant correlation between LL and lag-speaker-listener (lag-SL) couplings over time. Using the analogy of neural pattern (dis)similarity as distances between participants, we term this phenomenon the “herding effect”: like a shepherd guiding a group of sheep, the more closely listeners follow the speaker’s prior brain activity patterns (higher lag-SL similarity), the more tightly they cluster together (higher LL similarity). This herding effect is particularly pronounced in brain regions where neural synchronization among listeners tracks with behavioral ratings of narrative engagement, highlighting the mediating role of narrative content in the observed multi-brain neural coupling dynamics. By integrating LL and SL neural couplings, this study illustrates how unfolding stories shape a dynamic multi-brain functional network and how the configuration of this network may be associated with moment-by-moment efficacy of communication.

**Significance Statement:** Different stories have been found to evoke distinct brain activation patterns in the audience. This study delves into how the storyteller guides the audience through the multi-dimensional space of brain states, reflected in a series of shared activation patterns. We reveal that the listeners follow along the trajectory outlined by the speaker’s brain activity moments before, forming a tighter cluster at the more engaging moments of the story. This phenomenon is localized to high-level cortical areas supporting event representation. Our investigation illustrates how storytelling dynamically sculpts multi-brain neural dynamics in both the listeners and the speaker, shedding light on the potential association between the configuration of this network and communication efficacy.

## Introduction

Humans use narratives to convey complex, temporally structured sequences of thoughts to one another (Bruner et al., 1986; Willems et al., 2020). This kind of communication is thought to rely on a process of neural “alignment” or “coupling” (Pickering and Garrod, 2004; Hasson et al., 2012), whereby the speaker guides the listener(s) through a sequence of brain states to arrive at an understanding of the ideas or events the speaker intends to convey. Spoken stories have been found to drive synchronized neural alignment among listeners (LL coupling) throughout the cortical language network and extending into higher-level areas of the default-mode network thought to support event representation and narrative comprehension (Chen et al., 2016). On the other hand, asymmetric, time-lagged coupling has been observed between the speaker and listener(s) (SL coupling) in an overlapping set of high-level cortical areas (Stephens et al., 2010; Silbert et al., 2014; Zadbood et al., 2017; Liu et al., 2022; Nguyen et al., 2022; Zada et al., 2023).

The efficacy of a given narrative has been shown to vary across individuals. Both higher LL neural coupling and higher SL neural coupling have been separately associated with better behavioral estimates of speech comprehension across individuals (Stephens et al., 2010; Zadbood et al., 2017; Cohen et al., 2018; Zheng et al., 2018; Davidesco et al., 2019, 2019; Liu et al., 2019, 2022; Pan et al., 2020; Meshulam et al., 2021; Nguyen et al., 2022; Zhang et al., 2022, 2022; Zhu et al., 2022; Chen et al., 2023). Individuals performing better in the post-test often showed higher neural coupling with the speaker or other listeners. The efficacy of a narrative may also vary across time: a storyteller may meander or lose focus, and the content of the narrative may fluctuate in terms of how engaging it is or how much it resonates with listeners.

This study proposes a “herding hypothesis” incorporating both LL and SL neural coupling into a unified framework for the multi-brain neural dynamics between speaker and audience. Like a shepherd, a successful speaker guides the listeners toward the same brain states. We operationalize the “distance” between speaker and listeners as the intersubject (dis)similarity of brain activity patterns within different cortex regions: SL dissimilarity reflects the distance between the speaker and the listeners; LL dissimilarity reflects the distance between listeners (Fig. 1). The herding hypothesis proposes that when the listeners follow the speaker closely, they also tend to cluster more closely to each other. In other words, we expect SL and LL pattern (dis)similarities to correlate over the course of a narrative.

**Figure 1.**
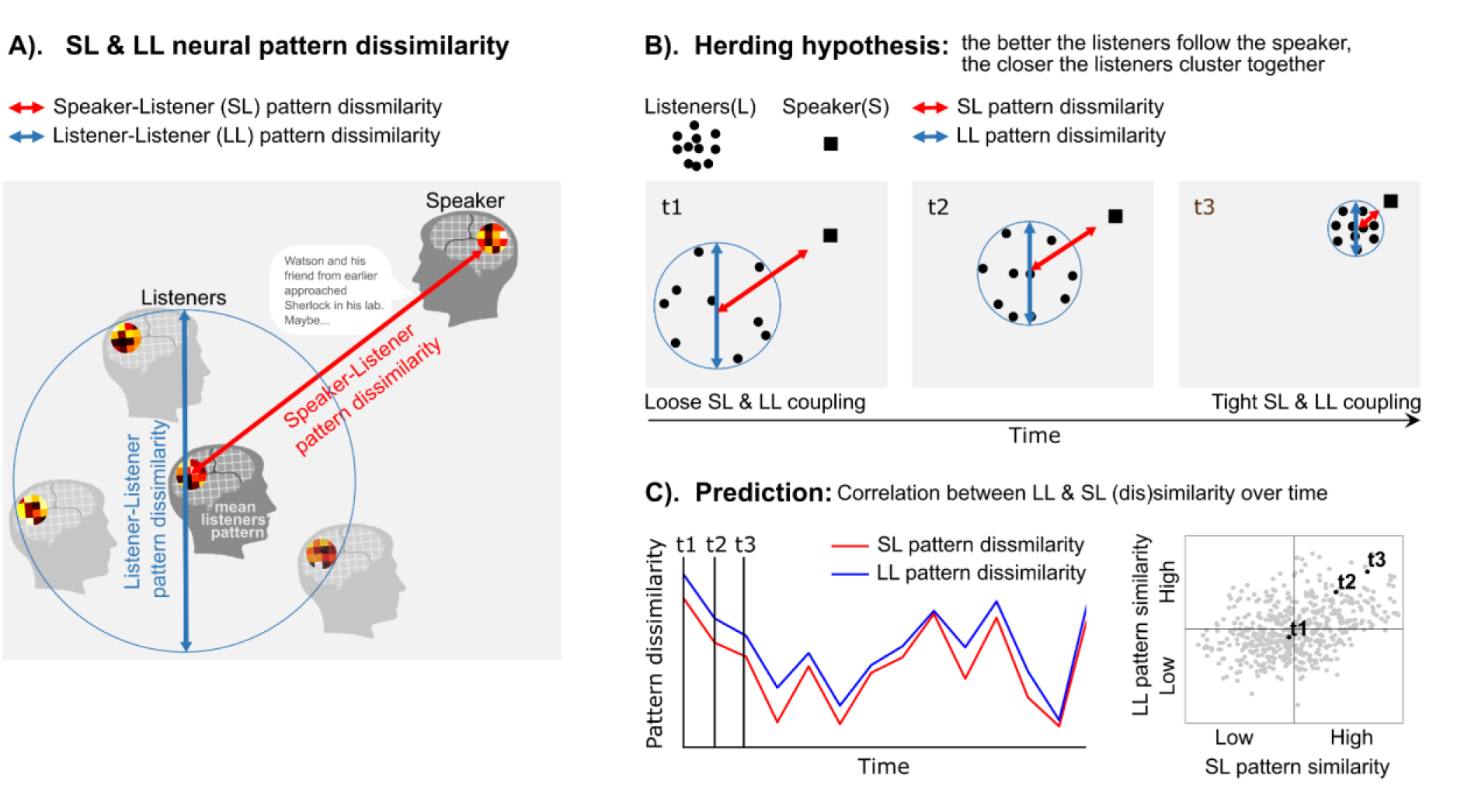
The herding hypothesis. A). We use neural pattern dissimilarities to quantify the distances between the speaker and listeners (SL) and the distances between listeners (LL). B). The herding hypothesis proposes that, over time, listeners will tend to cluster together when they follow the speaker closely, like a group of sheep guided by a shepherd. C). In other words, the herding hypothesis proposes that lag-SL pattern dis(similarity) will correlate with LL (dis)similarity over the course of a narrative. Note that SL coupling was computed using a lag of -10 to -1 TRs (speaker activity precedes for 15 to 1.5 seconds). See Extended Data Figure 1-1 for alternative hypotheses where SL and LL coupling converge and diverge in different ways.

In alignment with the shepherd analogy, we computed SL pattern dissimilarity with the speaker preceding the listeners across a window of lags ranging from -10 to -1 TR (1 TR = 1.5 seconds). The presence of a notable herding effect in this context would suggest that the listeners congregate around the trajectory outlined by the speaker’s brain activity 15 to 1.5 seconds earlier in time. Other plausible scenarios include listeners not following the speaker and dispersing in various directions (Finn et al., 2020), clustering together but deviating from the speaker’s path, or approaching the speaker but in a dispersed group (Extended Data Figure 1-1).

Aiming for a better understanding of the multi-brain neural dynamics underlying storytelling, we first verify the herding hypothesis: over time, the listeners tend to cluster together when they more closely follow the speaker and disperse in different directions when they deviate from the speaker’s neural trajectory. We then use a behavioral assessment of narrative engagement to illustrate that this herding effect is strongest for brain regions where listeners are most synchronized for the more compelling moments of the story.

## Materials and Methods

### fMRI datasets

This study relied on two openly available auditory story-listening datasets from the “Narratives” collection (OpenNeuro: https://openneuro.org/datasets/ds002245; (Nastase et al., 2021)), including “Sherlock” and “Merlin” (18 participants, 11 females). The speaker data reported in the original study (Zadbood et al., 2017) was also included. All participants reported fluency in English and were 18–40 years of age. The criteria for participant exclusion have been described in Zadbood et al. (2017). All participants provided informed, written consent, and the experimental protocol was approved by the institutional review board of Princeton University.

### fMRI preprocessing

fMRI data were preprocessed using FSL (https://fsl.fmrib.ox.ac.uk/), including slice time correction, volume registration, and high-pass filtering (140 s cutoff). All data were aligned to standard 3 × 3 × 4 mm Montreal Neurological Institute space (MNI152). A gray matter mask was applied. The first 25 and last 20 volumes of fMRI data were discarded to remove large signal fluctuations at the beginning and end of the time course to account for signal stabilization and stimulus onset/offset prior to computing intersubject dissimilarities (Nastase et al., 2019). The global mean responses were subtracted before pattern similarity analyses (Murphy et al., 2008; Garrido et al., 2013).

### ROI masks

We used 238 functional ROIs defined independently by Shen and colleagues (2013) based on whole-brain parcellation of resting-state fMRI data. ROIs with less than 10 voxels based on the coverage of our BOLD acquisition were excluded from further analyses.

### SL and LL neural similarities

We computed intersubject pattern correlations in each ROI at each time point of the story, i.e. TR by TR spatial pattern similarities (Fig. 1), using the leave-one-participant-out method (Nastase et al., 2019). For LL similarity, we computed the correlation between the activation pattern from one listener and the averaged pattern of the other 17 listeners. Similarly, SL similarity was computed between the speaker and the average pattern of 17 listeners, excluding each listener in turn. Note that quantifying SL coupling in this way entails that SL coupling can be high while LL coupling is low; i.e. listeners may be widely dispersed but roughly centered on the speaker. We also recomputed SL coupling by first computing the similarities between the speaker and each individual listener and then averaging these similarities. This analysis yielded qualitatively similar results (Extended Data Figure 3-5). According to the literature, the speaker and listener activation patterns are not necessarily temporally synchronized (Stephens et al., 2010; Dikker et al., 2014; Silbert et al., 2014; Zadbood et al., 2017; Liu et al., 2022). Therefore, we also computed the neural similarities at varying lags.

Pearson correlation was used to estimate pattern similarity. Time-lagged neural similarities were computed by circularly shifting the time series such that the non-overlapping edge of the shifted time series was concatenated to the beginning or end. The resulting correlation values were normalized with Fisher’s z transformation before further statistical analyses.

We statistically evaluated the SL and LL neural similarities separately before examining the herding effect (Fig. 2). We generated surrogates with the same mean and autocorrelation as the original time series by time-shifting and time-reversing the functional data prior to computing the intersubject similarities. We computed the correlation between the original seed and time-shifted/-reversed target time series. All possible time shifts were used to generate the null distribution. The resulting correlation values were compiled into null distributions after averaging across time points and participants. One-tailed z-tests were applied to compare neural similarities within the window of lag -10 to +10 TRs against this null distribution. We corrected for multiple comparisons across lags and ROIs by controlling the false discovery rate (FDR) at q < .05 (Benjamini and Hochberg, 1995).

**Figure 2.**
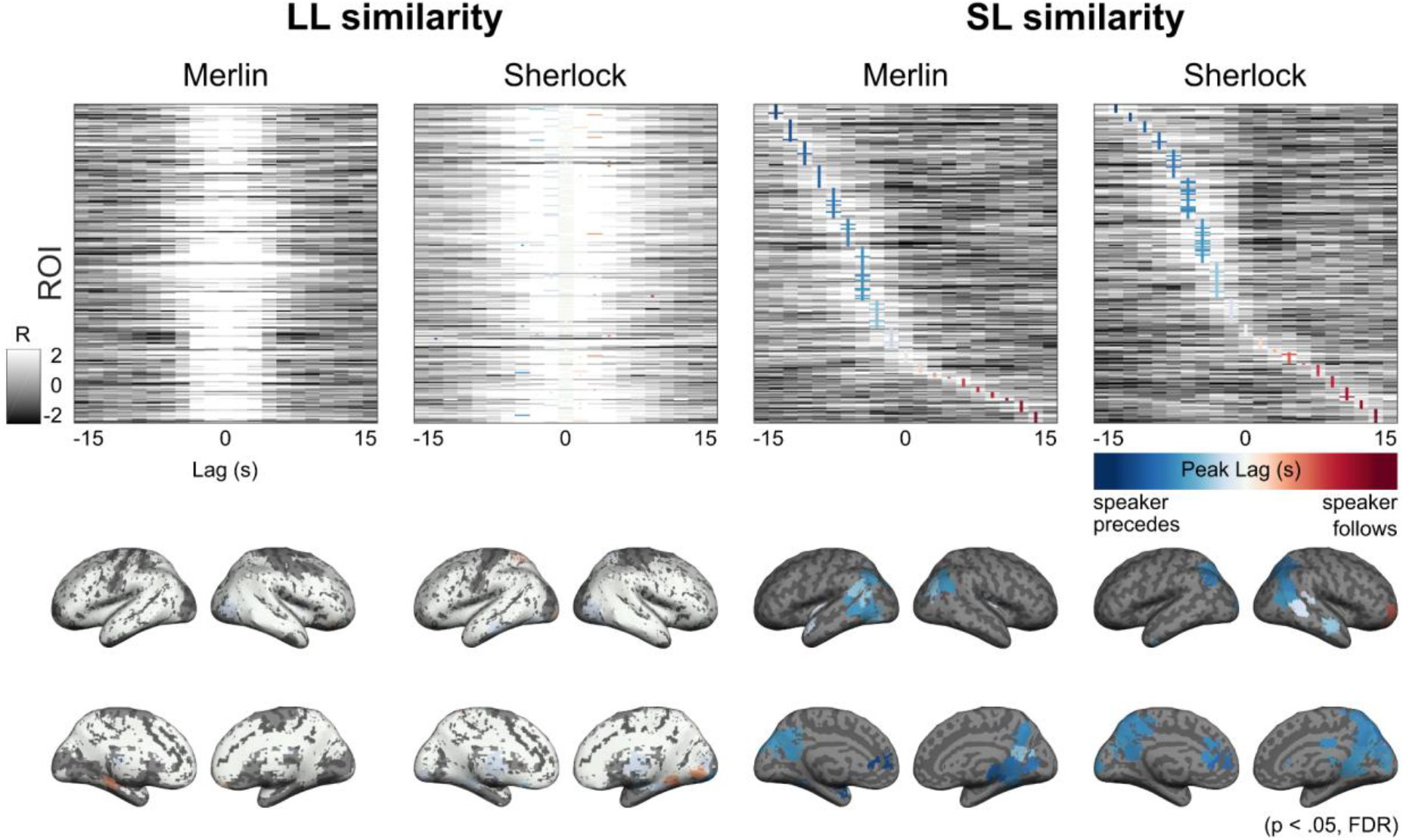
LL and SL neural pattern similarities. Full intersubject similarity matrices (upper) where each row shows the neural pattern similarities in each brain region at varying lags across columns. Brain regions are ordered by their peak lags. Intersubject pattern similarities for each TR were averaged across all TRs in the story. Lags with the peak correlation values are color-coded. Significant peak lags are marked with wide (horizontal) colored bars (p < .05, FDR correction). Nonsignificant peak lags are marked with narrow colored bars. The correlation values are normalized with Fisher’s transformation and then z-scored across lags. ROIs with significant peak lags are plotted on the brain (lower). LL similarities peak at lag 0 in most brain regions, reflecting that the listeners are synchronized. On the other hand, SL similarities often peak at negative lags, indicating that the speaker precedes the listeners.

### Computing the herding metric

We defined the herding metric as the correlation between LL neural similarity at lag 0 and SL neural similarity at lags within the window of -10 to -1 TRs (i.e. speaker precedes the listeners for 1.5 to 15 seconds). A significant herding effect indicates that the listeners are more synchronized when they echo the speaker’s activation pattern. Two statistical tests were applied.

First, to verify that only the speaker showed the herding effect, we replaced the actual speaker with each of the 18 listeners to serve as the pseudo-speaker (Extended Data Figure 3-1). LL similarity was computed among the remaining 17 listeners, excluding the pseudo-speaker, using the leave-one-out method, i.e. correlation between the activation pattern from one listener and the averaged pattern of all the other 16 listeners. SL similarity was computed between the real speaker and the average pattern of the 17 listeners, Pseudo-SL similarity was computed using the same method as the SL similarity, except that the real speaker was replaced by the pseudo-speaker. We computed the herding metric with the real and pseudo-SL similarity and compared the real and pseudo-herding effects using a two-sample one-tailed t-test (N = 18). We corrected for multiple comparisons across lags and ROIs by controlling the FDR at q < .05. Only the ROI x SL lag combinations that passed this test were included for the second statistical test.

Second, the speaker must precede the listeners to “herd” them. Therefore, we tested the real herding effect against correlation values between LL at lag 0 and SL at all the possible lags outside of the chosen lag window (-10∼-1 TR) using one-tailed z-tests. We circularly shifted the original time series to obtain a time-lagged time series. The number of possible lags equals the number of time points. The FDR method was used to control for multiple comparisons (ROI x SL lag; q < .05). Only ROIs that passed both statistical tests are considered to show a significant herding effect.

To quantify the amplitude of the herding effect, we extracted the peak LL-SL correlation value within the -10 to -1 TR SL lag window. We required that the peak value be larger than the absolute value of any negative peak and excluded any peaks occurring at the edges of the window.

### Behavioral engagement

#### Engagement ratings

Behavioral assessments of dynamic engagement were acquired in another group of participants recruited via Amazon Mechanical Turk. Participants with less than 20 unique rating scores (i.e. effectively flat ratings across the story) were excluded. 33 raters were included for “Merlin” (15 females). A separate sample of 34 raters was included for “Sherlock” (14 females). All participants reported fluency in English and were 25–71 years of age. All participants provided informed, written consent, and the experimental protocol was approved by the institutional review board of Princeton University.

The participants were instructed to indicate “how engaging the current event is” while listening to the stories by moving a slider continuously. We presented the stories and collected the data using the web-based tool DANTE (Dimensional Annotation Tool for Emotions) ( https://github.com/phuselab/DANTE) (Boccignone et al., 2017). The rating scores were acquired with a resolution of 0.04 seconds and then downsampled to 1.5 seconds (= 1 TR).

The engagement scores were z-scored across time, detrended, and averaged across raters.

#### Correlation between engagement and LL neural similarity

To quantify the relationship between time-point-by-time-point engagement ratings and LL neural similarity, we computed the Pearson correlation between the engagement scores and the LL similarity over time within ROIs showing a significant herding effect (one-tailed). We corrected for multiple comparisons across ROIs by controlling the FDR at q < .05.

#### Correlation between engagement effect and herding effect across ROIs

To quantify the relationship between engagement ratings and group-level herding, we computed the Pearson correlation between the herding effect and the engagement effect across all ROIs (p < .05). Note that since the number of ROIs is fixed, a significant p-value associated with this correlation does not indicate generalization to other regions (and does not support population-level inference).

## Results

The herding hypothesis predicts that listeners will more closely cluster together at moments of the story where they more closely follow the speaker (Fig. 1). We quantify the distance between speaker and listeners by computing the moment-by-moment intersubject (dis)similarity of neural activity patterns. The resulting dynamic LL and SL couplings indicate how tightly the listeners are clustered together and aligned to the speaker, respectively. We calculate the SL dissimilarity at different lags from -15 to -1.5 seconds. We first verify that listener activity patterns echo those of the speaker with a lag of several seconds; that is, SL similarities peak at negative lags (Fig. 2). We then reveal a significant herding effect in which the LL coupling is correlated with the strength of lag-SL coupling in the DMN and language network (Fig. 3). Finally, we show higher LL coupling at moments of the story behaviorally reported as more engaging (Fig. 4A). This effect is stronger in brain regions showing higher herding effect, such as DMN (Fig. 4B), providing behavioral evidence that the herding effect reflects how well the audience follows the speaker.

**Figure 3.**
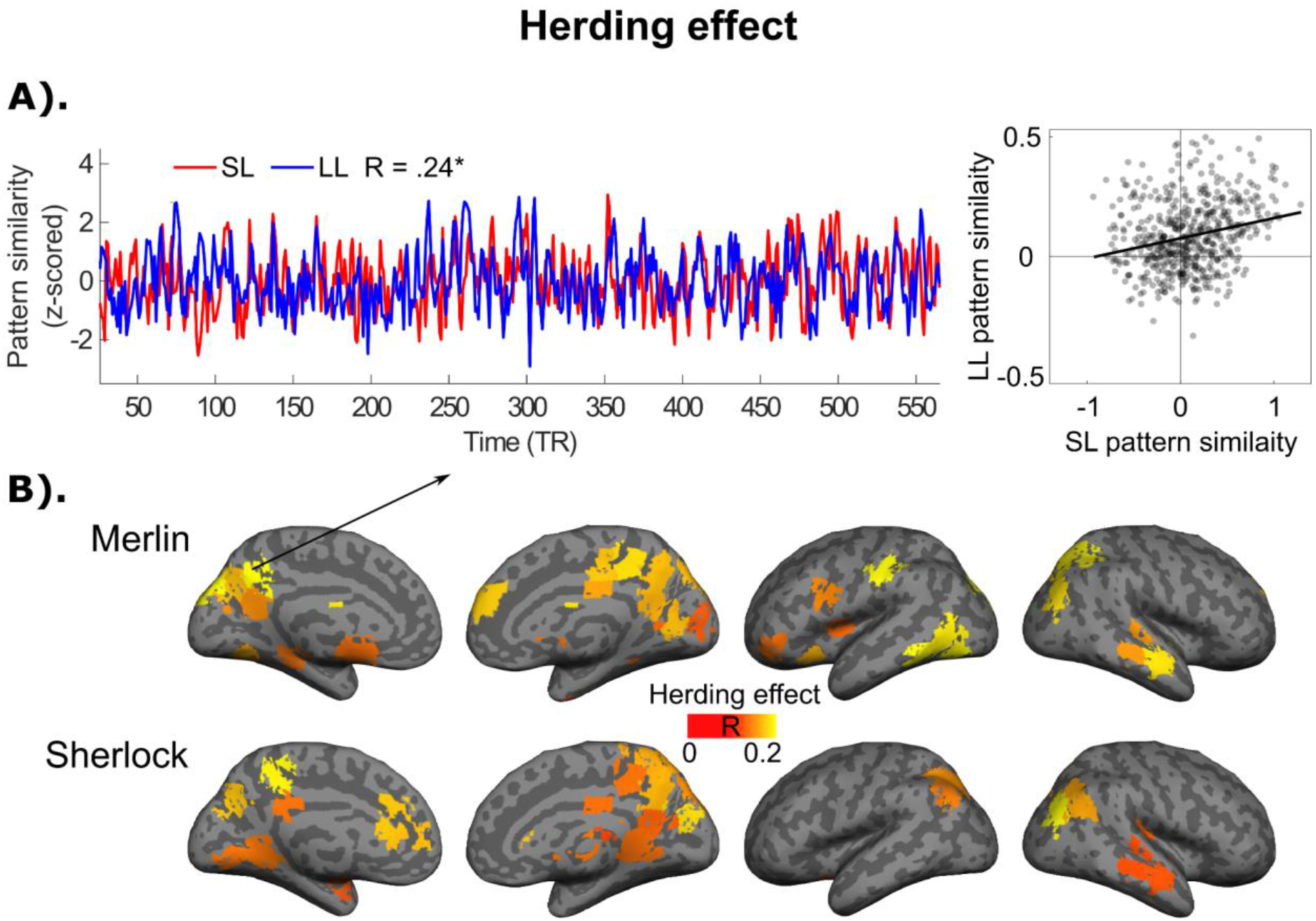
Cortical areas with a significant herding effect. A). A left precuneus ROI showing a significant correlation between lag-SL and LL coupling over the course of “Merlin,” namely, a significant herding effect. B). All ROIs with a significant herding effect. They are colored according to the amplitude of the correlation between lag-SL and LL similarity (p < .05, FDR correction). lag-SL and LL similarities were normalized with Fisher’s transformation and z-scored across time before computing the herding effect.

**Figure 4.**
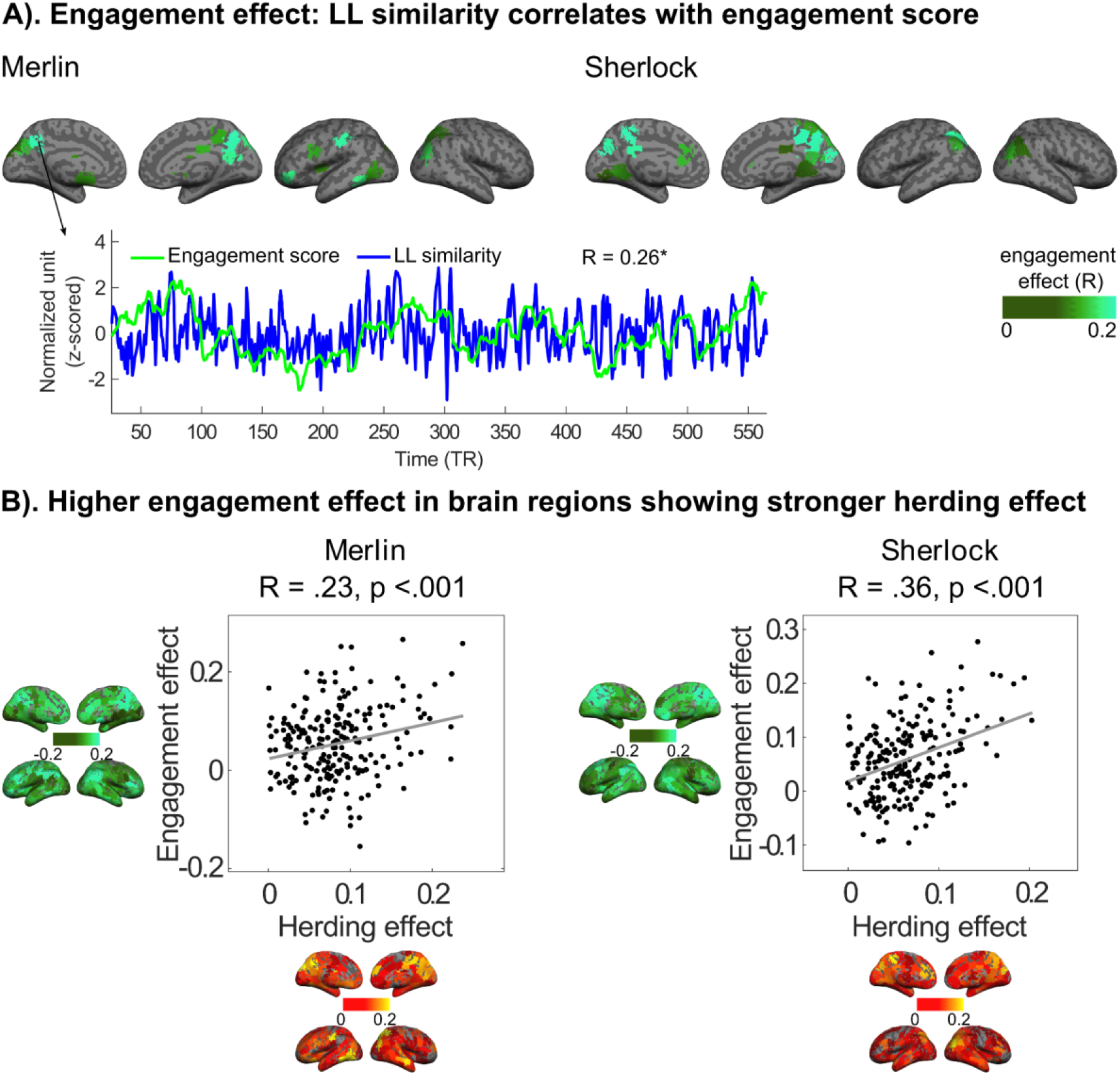
The engagement effect. A). Brain regions where the behavioral engagement score significantly correlated with LL neural similarity (p < .05, FDR correction). In these regions, moments in the story with higher engagement ratings elicit higher LL neural similarity. The color bar indicates the magnitude of the correlation between the engagement score and LL similarity. B). Brain regions with a stronger herding effect tend to show a stronger engagement effect. This finding provides behavioral evidence that the herding effect reflects how engaging listeners find the narrative.

### SL and LL neural similarities: Listeners follow the speaker’s brain activity patterns

To visualize the relationship between SL and LL neural similarities, we first plot the total ROI × lag intersubject similarity matrices (Fig. 2). In agreement with previous studies (Stephens et al., 2010; Dikker et al., 2014; Zadbood et al., 2017; Liu et al., 2022), SL dynamics are markedly different from LL dynamics. LL similarities peak at lag 0 in most regions: listener brain activity patterns are synchronized in processing the story’s content. In contrast, significant SL similarities mainly occur at negative lags: listener activity patterns echo the speaker activity patterns with seconds-long lags.

### Herding effect: The more closely the listeners follow the speaker, the more tightly the listeners cluster together

The herding hypothesis predicts that the more closely the listeners follow the speaker, the more closely the listeners cluster together. We quantified the herding effect by computing the correlation between moment-by-moment LL coupling and lagged SL coupling throughout the narrative. Statistical significance for the herding effect was assessed using permutation procedures based on two surrogate datasets, one generated by replacing the speaker with a “pseudo-speaker” sampled from the listeners (Extended Data Figure 3-1), the other by applying unreasonable SL lags (i.e., the speaker precedes the listeners for more than 15 seconds or the speaker does not precede the listeners). Only ROIs that passed both statistical tests are considered to show a significant herding effect. Namely, a stronger herding effect was found with the real speaker rather than pseudo-speakers and with reasonable rather than unreasonable SL lags. For comparison, we also computed the herding metrics based on SL coupling at 0 lag and found no significant effect (Extended Data Figure 3-2). We test the herding hypothesis with one-tailed tests. For the results of ad hoc two-tailed tests, see Extended Data Figure 3-3.

Our results reveal a significant herding effect for both stories in the precuneus, posterior cingulate cortex, cuneus, superior and middle temporal gyrus, and superior/middle occipital gyrus (Fig. 3); many of these regions have been implicated in representing high-level events and narrative features (Chen et al., 2016; Baldassano et al., 2017, 2018; Chang et al., 2021). See Extended Data Figure 3-4 for an exemplar ROI showing significant LL coupling but a nonsignificant herding effect. In addition, a similar herding effect is revealed with alternative SL coupling measurement, namely, averaged pattern similarity between the speaker and each listener (Extended Data Figure 3-5) instead of pattern similarly between the speaker and the averaged listener pattern (Fig. 3).

### More “herded” brain regions show stronger LL similarity at engaging moments of the story

Narratives serve as the conduit for the speaker’s information to reach the listeners. We hypothesized that fluctuations in how engaging listeners find certain parts of the story may relate to how effectively the speaker “herds” the listeners. To behaviorally assess how engaging the spoken narrative was moment by moment, we collected continuous engagement scores from a separate group of participants. In agreement with a previous study (Song et al., 2021), we found that engagement scores correlate with LL neural similarity; that is, higher LL similarity occurs at more engaging moments of a story. In a similar vein, higher neural synchronization has been reported for more memorable (Simony et al., 2016), surprising (Brandman et al., 2021), and emotional moments during stories (Nummenmaa et al., 2014; Smirnov et al., 2019).

Among regions showing a significant herding effect, a significant engagement effect was found for both stories in the precuneus, posterior cingulate cortex, cuneus, and superior/middle occipital gyrus (Fig. 4A). More importantly, the engagement effect is larger in areas showing a stronger herding effect (Fig. 4B). This finding provides behavioral evidence that the herding effect reflects how engaged the listeners are with the content of the story.

## Discussion

This study examined the multi-brain neural dynamics underlying storytelling. We first verified that the audience echoed the speaker’s neural activation patterns with a temporal lag (Fig. 2) (Stephens et al., 2010; Dikker et al., 2014; Zadbood et al., 2017; Zheng et al., 2018; Davidesco et al., 2019; Liu et al., 2022). As predicted by the herding hypothesis (Fig. 1), the more closely the listeners’ brain activity matched that of the speaker, the more closely the listeners clustered together (Fig. 3). We argue that this herding effect is an index of effective communication during storytelling, indicating that, metaphorically, the speaker guides the listeners’ neural activity, especially in higher-order brain areas. We also show that LL neural similarity increases at more engaging moments of the story (Fig. 4A). This engagement effect is stronger in the more “herded” brain regions (Fig. 4B), supporting the hypothesis that the herding effect reflects effective storytelling—that is, when the storyteller most successfully conveys their thoughts to the listeners.

A significant herding effect was found in several high-order brain areas in the DMN, including the precuneus, middle/posterior cingulate cortex, lateral parietal cortex, and right anterolateral temporal cortex (Fig. 3). The posterior medial regions in particular have been shown to encode paragraph-level narrative structure (Lerner et al., 2011). These regions are thought to host content-specific, supramodal event representations (Honey et al., 2012; Chen et al., 2016; Baldassano et al., 2017; Yeshurun et al., 2017; Nguyen et al., 2019; Chang et al., 2021), linking the production and comprehension of spoken narratives (Chen et al., 2017; Zadbood et al., 2017; Liu et al., 2022). The current results add a dynamical perspective to this body of work, suggesting that the speaker’s own neural trajectory through high-level event features may guide the listeners’ upcoming event representations with varying effectiveness over the course of a narrative. This dynamic convergence and divergence of idiosyncratic internal representations with the unfolding narrative would be particularly interesting for further investigation.

Note that we do not observe a significant herding effect with SL at lag 0 (Extended Data Figure 3-2), indicating that the speaker and listeners are not simultaneously synchronized by low-level auditory features of the story. That is, significant herding effect emerges at negative lags on the scale of several seconds (6 seconds on average, ranging from 3 to 12 seconds across ROIs and stories), and cannot be accounted for by low-level stimulus features alone. The scale of these lags is consistent with work demonstrating that higher-level narrative features and event-level representations are constructed over several seconds along the cortical processing hierarchy when listening to naturalistic narratives (Chang et al., 2022). The pseudo-speaker analyses (Extended Data Figure 3-1)—i. e. systematically permuting the speaker with each listener in turn—also affirm that the herding effect is not a result of shared stimuli. Instead, the speaker *leads* the listeners through a trajectory of brain states over the course of the narrative. Consequently, the listeners tend to cluster along the speaker’s path, displaying varying degrees of proximity with each other—sometimes tightly, sometimes loosely—but consistently remaining a few steps behind the speaker.

If the herding effect can be interpreted as an index of effective speech, what factors may impact how closely the speaker resonates with the audience? In keeping with recent theoretical work positioning the DMN as a high-level interface between external events with prior knowledge (Yeshurun et al., 2021), we speculate that individual differences in how closely listeners follow the speaker may in part reflect differences in the way the speaker’s narrative aligns with each listener’s internal state and idiosyncratic memories. Prior work, for example, has suggested that brain-to-brain coupling may vary as a function of social closeness (Dikker et al., 2017; Bevilacqua et al., 2019) or whether the speaker and listener share similar beliefs (e.g. similar political orientation (Katabi et al., 2023)). It is worth noting, however, that there may be different kinds of effective speech. For example, a speaker may seek to (mis)direct listeners toward a different understanding than their own; listeners that are unfamiliar versus experts with a particular topic may experience the same speech very differently (Meshulam et al., 2021; Chen et al., 2023); and a speaker may attempt to “meet certain listeners where they are” rather than wrangling all listeners similarly.

The methodology we introduce takes into account both LL and SL couplings in a moment-by-moment manner. In our stories, most of the time, the listeners converge to trail the speaker and when they lose track, they disperse in different directions (low LL and low lag-SL; time points in the lower-left quadrant of the scatterplot in Fig. 3A). However, there are moments where LL is high despite low lag-SL (time points in the upper-left quadrant of the scatterplot in Fig. 3A, also see Extended Data Figure 1-1), or LL is low despite relatively high lag-SL (e.g. time points in the lower-right quadrant of the scatterplot in Fig. 3A). We speculate that in the former case, the audience might share the same misunderstanding, or the speaker might not undergo the experience from the same perspective as the listeners (Sun et al., 2020) while in the latter case, the listeners might only form a loose group around the speaker due to ambiguous speech or heterogeneous apprehension (Nguyen et al., 2019).

We hope that, with more diverse speakers and larger audiences, future work will be able to more extensively sample the less successful moments of communication across narratives. Our framework for measuring dynamic, multi-brain coupling can highlight when and how a speaker and the audience become misaligned. By relating these moments back to the speaker’s delivery, narrative content, and the personal backgrounds of both the speaker and individual listeners, we will be better positioned to identify why communication breaks down and develop solutions for accommodating different learning styles.

## Data and code availability

This study relied on openly available spoken story datasets from the “Narratives” collection (OpenNeuro: https://openneuro.org/datasets/ds002245) (Nastase et al., 2021).

## Acknowledgments

This study is supported by the National Institute of Mental Health (R01-MH112357 and DP1-HD091948).

## Author contributions

C.H.C.C. designed research; C.H.C.C. performed research; C.H.C.C. contributed new reagents/analytic tools; C.H.C.C. analyzed data; and C.H.C.C., S.A.N., and U.H. wrote the paper.

