## Extended Data for "How a speaker herds the audience: Multi-brain neural convergence over time during naturalistic storytelling"

### Supplemental information

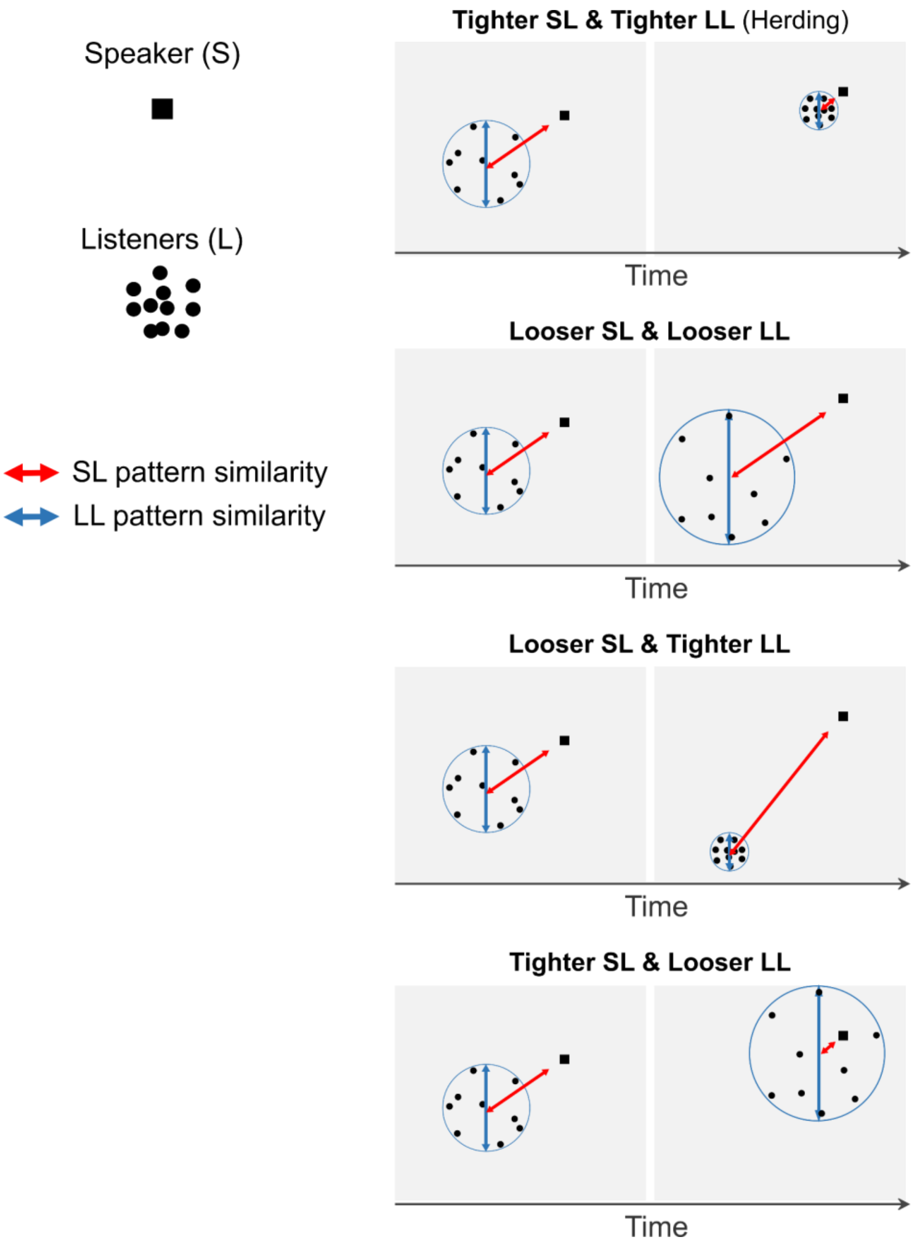


**Extended Data Figure 1-1**. Alternative scenarios to the herding hypothesis. While the herding hypothesis suggests that, over time, listeners will tend to tightly cluster when closely following the speaker, alternative situations may arise. These include listeners not following the speaker and dispersing in various directions, clustering together but deviating from the speaker's path, or approaching the speaker as a dispersed group. Note that SL coupling was computed using a lag of -10 to -1 TRs.

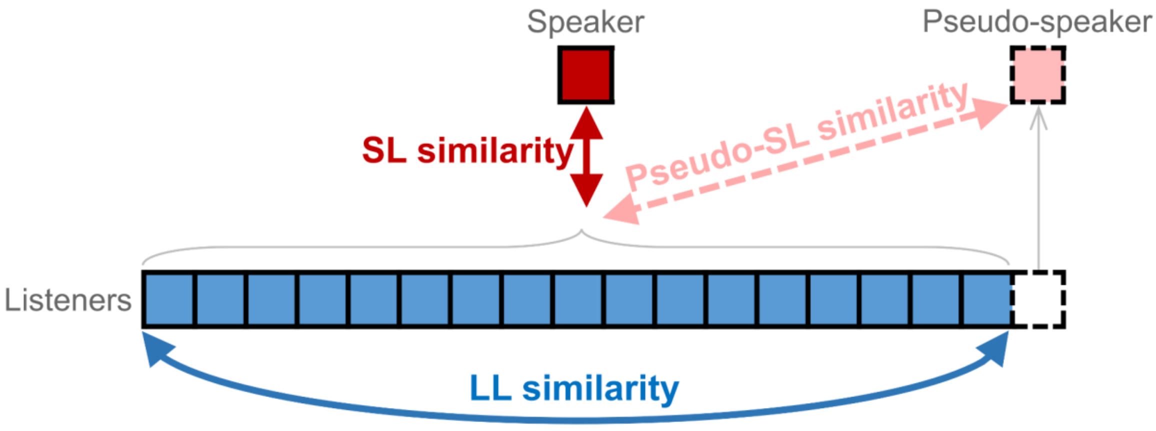


**Extended Data Figure 3-1**. To verify that only the speaker shows the herding effect, we took each listener as the pseudo-speaker in turn. The real herding effect was tested against the correlation values between LL similarities and pseudo-SL similarities.

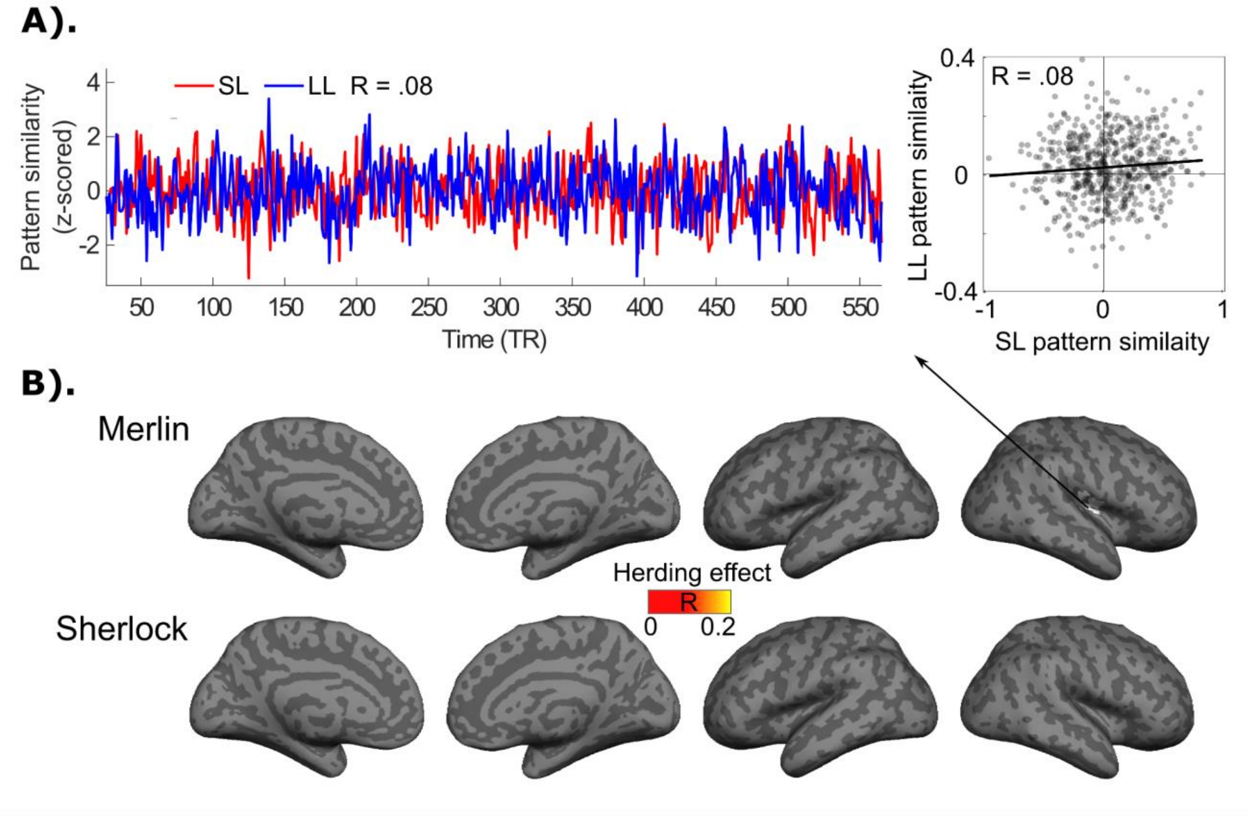


**Extended Data Figure 3-2.** Herding metrics computed with SL coupling at 0 lag. A). A right rolandic operculum/insula ROI showing a nonsignificant herding effect despite significant LL coupling. B). None of the ROIs show a significant herding effect in either story with SL coupling at lag 0 ($p < .05$, FDR correction).

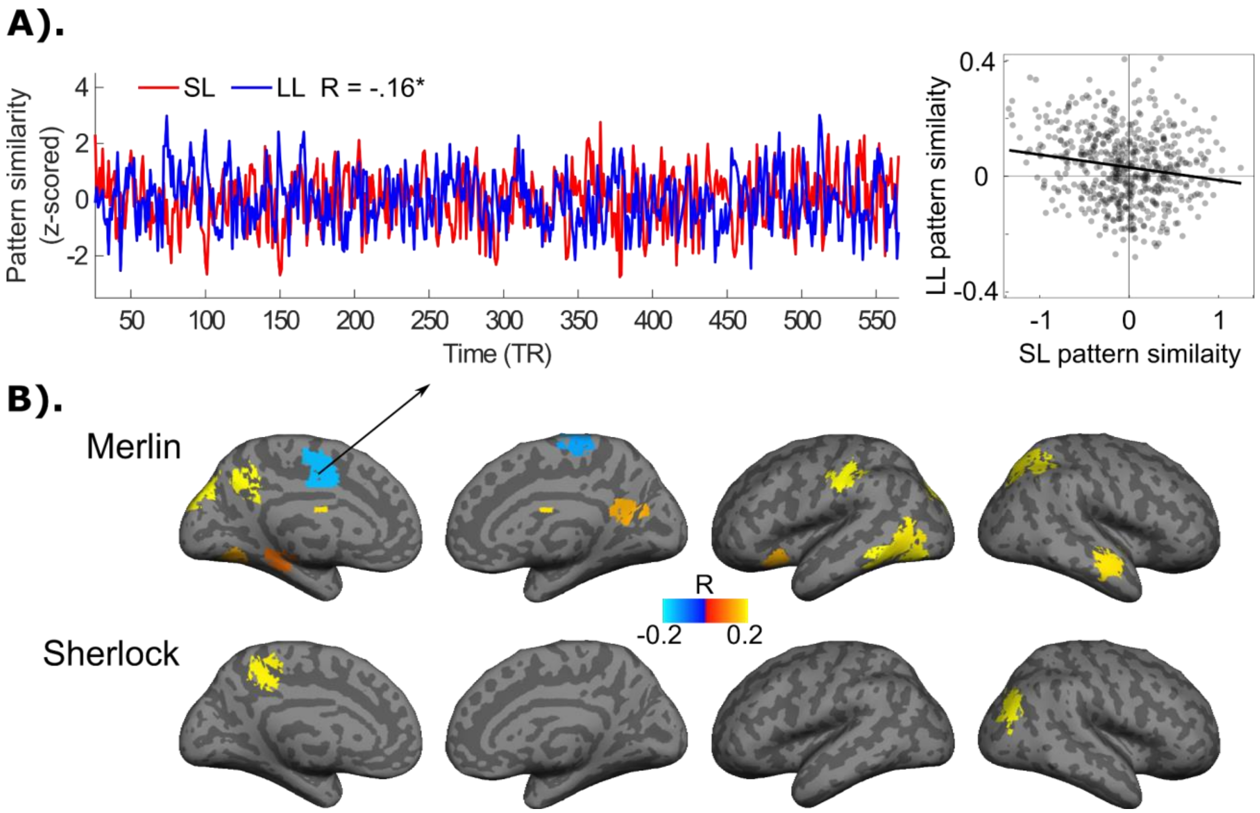


**Extended Data Figure 3-3. Herding metrics under ad hoc two-tailed tests.** A). A left supplementary motor area ROI showing a negative correlation between lag-SL and LL coupling. B). All ROIs with a significant non-directional correlation between lag-SL and LL coupling ($p < .05$, FDR correction). See Fig. 3 for the herding metrics under one-tailed tests.

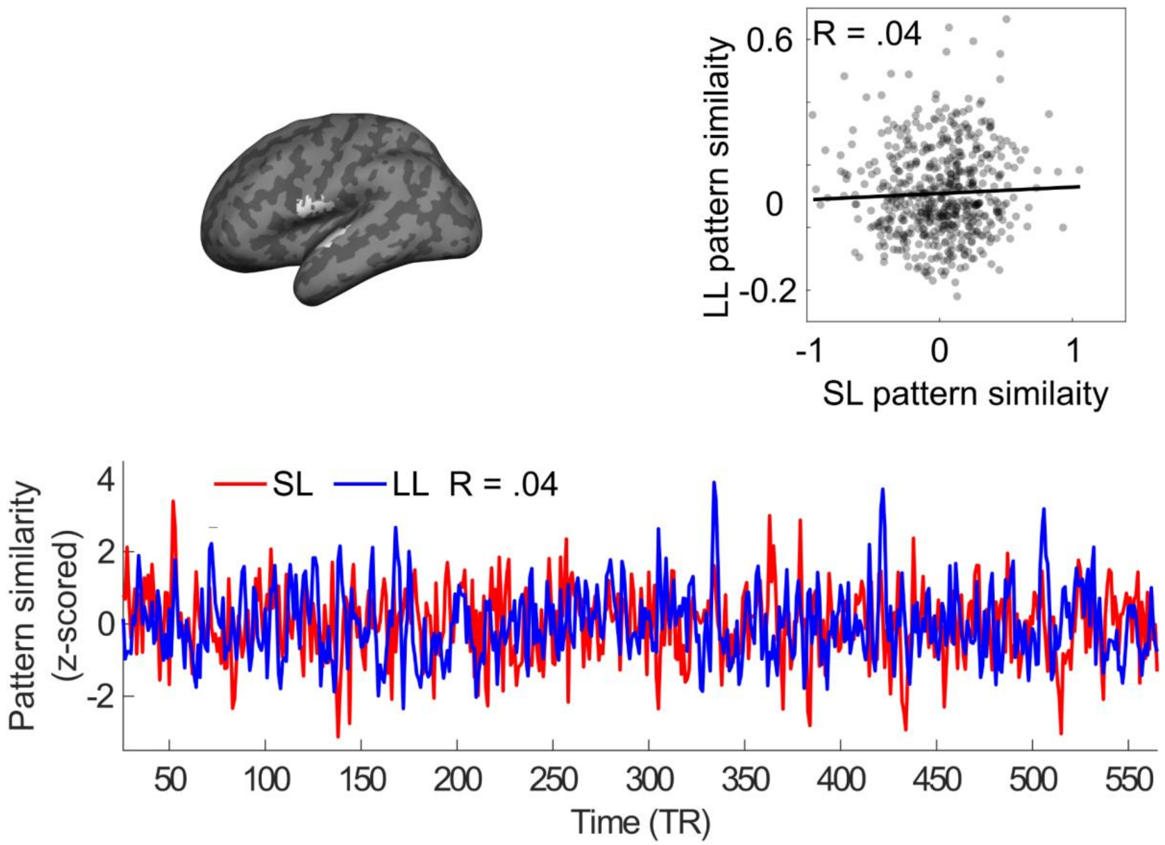


**Extended Data Figure 3-4.** An example ROI showing a nonsignificant herding effect. Left superior temporal gyrus and precentral/postcentral gyrus ROI showing a nonsignificant herding effect ($R = .04$, $p > .05$, FDR correction), but significant LL coupling ($R = .11$, $p < .05$, FDR correction) in "Merlin."

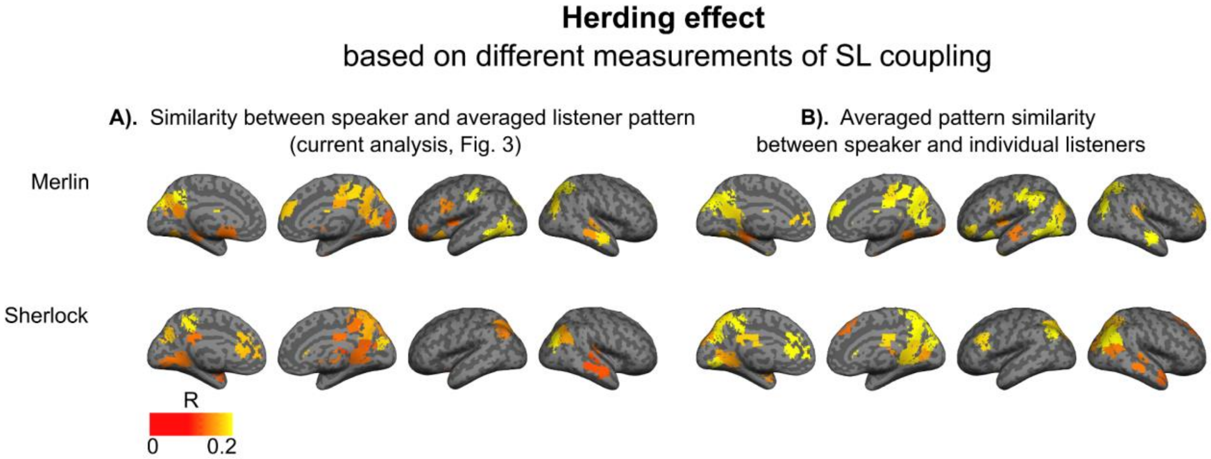


**Extended Data Figure 3-5.** Herding effect based on different measurements of SL coupling. We quantified lag-SL coupling in two ways: A). computing the similarity between the speaker activity pattern and the average listener pattern; or B). computing the similarities between the speaker and each individual listener, then averaging those similarities. ($p < .05$, FDR correction)
